# Adaptive genomic variation is linked to a climatic gradient in a social wasp

**DOI:** 10.1101/2023.10.12.561994

**Authors:** Hannah L. Cook, Sara E. Miller, Gilbert Giri, Kevin J. Loope, Michael J. Sheehan, Floria M.K. Uy

**Affiliations:** Department of Biology, University of Rochester, NY, USA; Department of Biology, University of Missouri–St. Louis, St Louis, USA; Department of Fish and Wildlife Conservation, Virginia Tech, Virginia, USA; Laboratory for Animal Social Evolution and Recognition, Department of Neurobiology and Behavior, Cornell University, NY, USA; Department of Entomology, Washington State University, WA, USA

**Keywords:** climatic gradient, genomic variation, genotype-environment associations, local adaptation, *Mischocyttarus mexicanus*, social wasp

## Abstract

Species vary in their ability to adapt to rapid changes, with the presence of genetic variation often facilitating long-term evolutionary responses. Given the impending threat of climate change, it is critical to investigate how genetic variation facilitates persistence and possible range expansion in animals. Here, we combine genomic and climatic data to characterize the drivers of local adaptation in the widely distributed, social wasp *Mischocyttarus mexicanus cubicola*. Using whole genome sequence data, we show that adaptive genomic variation is linked to a climatic gradient across the broad distribution of this species. We found strong population structure, dividing the species into two genetic clusters that follow subtropical and temperate regions. Patterns of gene flow across the range deviate from those expected by isolation by distance alone with climatic differences resulting in reduced gene flow even between adjacent populations. Importantly, genotype-environment analyses reveal candidate single nucleotide polymorphism (SNPs) associated with temperature and rainfall, suggesting adaptation for thermal and desiccation tolerance. In particular, candidate SNPs in or near mitochondrial genes *ND5*, *CO1*, and *COIII* are linked to cold tolerance and metabolism. Similarly, the *Gld* nuclear gene shown to mediate cold hardiness and cuticle formation, shows two candidate SNPs with non-synonymous mutations unique to temperate populations. Together, our results reveal candidate SNPs consistent with local adaptation to distinct climatic conditions. Thus, the integration of genomic and climatic data can be a powerful approach to predict vulnerability and persistence of species under rapid climate change.

## 1 INTRODUCTION

Some species show remarkable abilities to adjust to environmental changes, which can be mediated by phenotypic plasticity and/or genetically-based adaptations (Nicotra et al. 2015, Thompson et al. 2023). Understanding how selective and demographic forces shape this adaptive potential is critical to elucidate adaptive responses and, ultimately, population resilience (Holderegger et al. 2006, Balkenhol et al. 2019). To this end, the distribution of genomic variation across the range of a widely distributed species can provide insights into the complex interactions between the genome and the variable environment that ultimately facilitate adaptation (Hoffmann et al. 2021). For instance, extensive genetic variation increases the probability that a population could respond adaptively to changing local conditions (Hughes et al. 2008). In addition, neutral genetic markers facilitate characterization of gene flow, population structure and dispersal, thereby providing insight into the ability of a species to colonize and persist in diverse environments (Holderegger et al. 2006). Therefore, species that occupy environmental gradients provide the unique opportunity to detect patterns of adaptive variation of local populations exposed to distinct or changing climatic pressures (Balkenhol et al. 2019, Hofmann et al. 2021).

Under the imminent threat of rapid climate change, elucidating how bioclimatic variables interact with genetic variation is critical in predicting if and how species will persist (Franks and Hoffmann 2012). Recent genome-wide studies found that temperature and precipitation are important selective pressures in animals with broad distributions. For instance, latitudinal clines of house mice (*Mus musculus domesticus*) along the West and East Coast of North America show convergence in patterns of body size evolution, facilitated by parallel genetic changes in response to similar climatic gradients (Ferris et al. 2021). Genetic variation in European starlings (*Sturnus vulgaris*) is also linked to changes in temperature and precipitation across their broad range where introduced in North America (Hofsmeister et al. 2021). Studies in ectothermic animals provide similar evidence of response to variable habitats in genes known to mediate temperature and precipitation tolerance. For example, a study in natural populations of *Drosophila melanogaster* across Europe and North America identified candidate single-nucleotide polymorphism (SNPs) linked to variation in temperature, rainfall, and wind (Bogaerts-Márquez et al. 2021). These two independent, broad geographical distributions also showed parallel signatures of seasonal adaptation across similar climatic gradients (Machado et al. 2021). Two species of montane bumblebees also show variation in thermal and precipitation tolerance, showing evidence for selection on genes that mediate facultative endothermy and desiccation tolerance (Jackson et al. 2020, Heraghty et al. 2022). However, few studies have effectively characterized local adaptation by investigating genome-wide patterns of selection and responses of traits specifically associated with geographic and climatic variation (see Samad-zada et al. 2023).

Most studies exploring adaptations to climate focus on physiological and morphological traits. However, climate is also a major driving force in the evolution of sociality, mediating social adaptive responses such as grouping behavior (Johnson et al. 2023, Ostwald et al. 2023). For example, harsh climatic conditions drove the evolution of sociality in Australian rodents (Firman et al. 2020). In the greater ani (*Crotophaga major),* the size of cooperative breeding groups is influenced by fluctuating climate (Riehl & Smart, 2022). Many bees also adjust their level of sociality depending on environmental conditions (Wcislo and Fewell 2017), including forming groups to conserve body mass and heat in the winter (Ostwald et al. 2022). Similarly, Polistine social wasps distributed globally and exposed to different climates, vary in their nesting behavior (Miller et al. 2018, Sheehan et al. 2015) and thermoregulatory responses (Stabentheiner et al. 2022). Comparative analyses of genomic variation across populations therefore can identify candidate genes involved in shaping local adaptation and facilitate predicting future ranges and sociality in cooperative species (Perez and Aron 2020, Hartke et al. 2021).

Here, we characterize the main drivers of local adaptation for the social wasp, *Mischocyttarus mexicanus cubicola*, a species with a broad distribution throughout the Southeast and the gulf coasts of the United States, with a recent expansion to Texas (Carpenter et al. 2009, Kratzer 2022; Fig 1A). In the Southeast, this species ranges from South Florida to South Carolina encompassing a broad latitudinal and climatic range (Hermann and Chao 1984b; Fig. 1A). The range spans the transition between subtropical and temperate zones, and this transition is associated with differences in nest initiation synchrony, colony formation and behavior (Hermann and Chao 1984a, Clouse 1995). For instance, subtropical populations are active throughout the year while temperate populations have adapted to a seasonal climate and exhibit limited activity during winters (Litte 1977, Gunnels IV 2007, Mora-Kepfer 2011). Further, nest building is asynchronous and occurs year-round in the subtropics (Mora-Kepfer 2014), but is limited to the late spring and summer in temperate zones (Litte 1977, Gunnels IV 2007).

**Fig 1.**
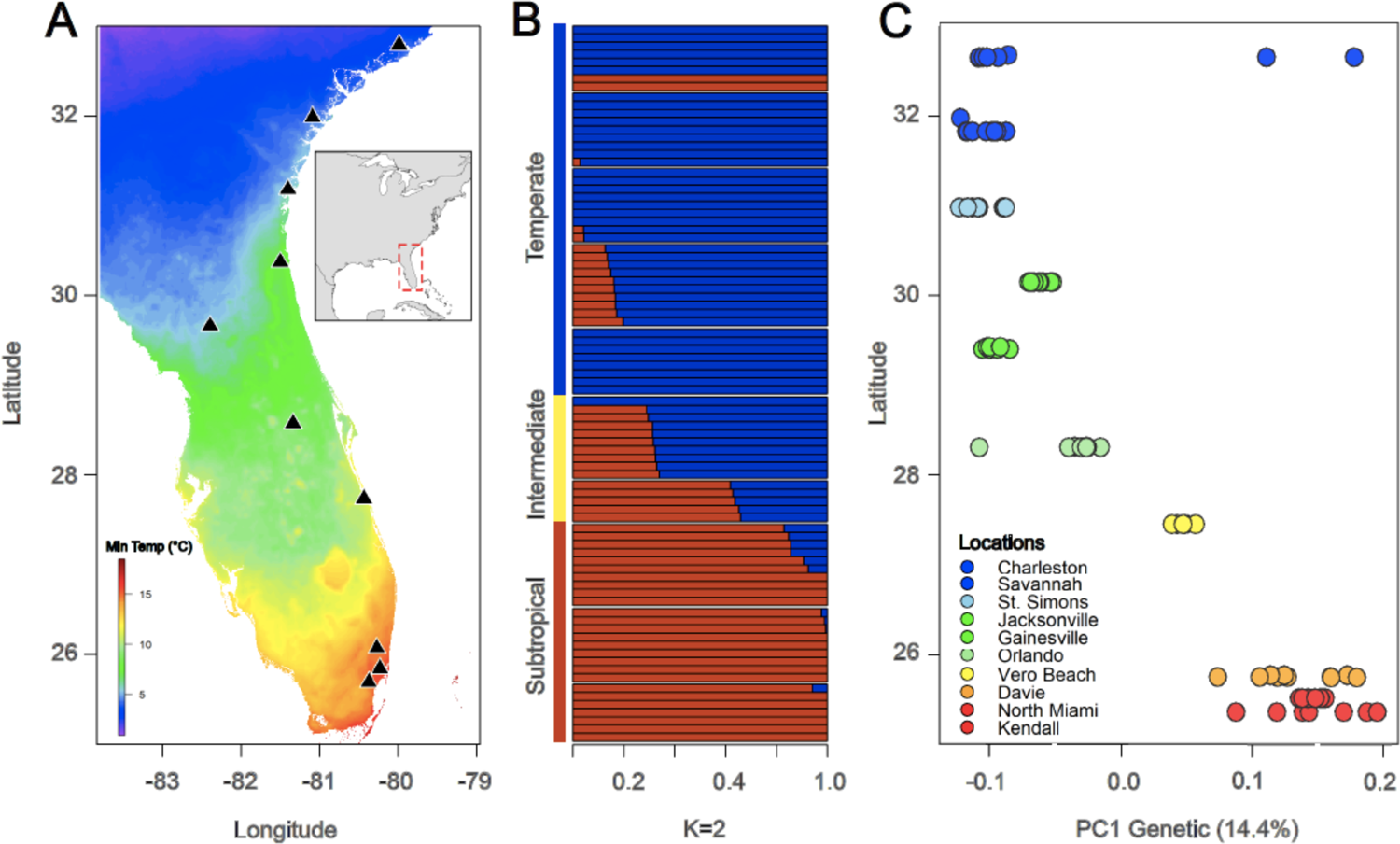
Population structure and genetic variation across the climatic gradient. **A.** The transect map depicts the mean daily minimum temperature during the coldest month in °C (BIO6) for the ten sampling sites across the transition from subtropical to temperate zones. **B.** Ancestry admixture plot highlights two genetic clusters that follow the: subtropical (red) and temperate (blue) zones (K =2). The transition zone (yellow) shows populations with mixed ancestries. **C.** PC1 Genetic explains 16% of the genetic variation and follows the climatic pattern across the latitudinal transect, separating southern from northern populations and showing the same intermediate pattern for the transition zone (yellow). Two northern outliers group with the southern genetic cluster.

Changes in group formation and behavior also follow this gradient. Subtropical populations form foundress groups that often accept non-nestmates to initiate cooperative nest building (Mora-Kepfer 2014). However, populations in the transition and temperate zones show a higher incidence of single foundresses that build nests in the spring, and a higher tolerance for forming groups in the summer and early fall (Litte 1976, Gunnels et al. 2008).

Grounded on this known link between climate and variation in nesting phenology and colony formation strategies across the *M. mexicanus* range, we predicted that population genomic patterns might also follow this latitudinal and climatic cline. These patterns should reflect both demographic and adaptive processes mediating local adaptation and range expansion. Specifically, we first predict that gene flow and population structure will be mediated both by distance between populations and by climatic boundaries. That is, gene flow should be most apparent between adjacent populations, but will be reduced across the transition from tropical to temperate climate. Second, we predict signatures of local adaptation associated with the climatic gradient, especially in genes linked to thermal tolerance, desiccation, and diapause. For instance, spatial distribution of specific genotypes will be associated with changes in the minimum temperature during winter and/or amount of precipitation over summer, which vary dramatically between subtropical and temperate regions. Genes facilitating tolerance for temperature and drought should be affected. Together, our work disentangles the relative effects of demography, geography and local climate in shaping genomic variation, providing insights on the mechanisms underlying range expansion of a widely-distributed social insect.

## 2 MATERIALS AND METHODS

### 2.1 Data acquisition

During 2018 and 2019, we collected samples for 10 populations of *M. mexicanus* across the Southeastern USA (Fig 1A). Our latitudinal transect consisted of the transition from the subtropical to temperate zones. Altitudes ranged from 3 to 54 meters above sea level, which removed elevational effects. We collected from three subtropical populations in South Florida, two populations located in the transition from subtropical to temperate climate in Central Florida, and six temperate populations starting in North Florida with the northernmost population located in South Carolina (Fig 1A). We collected one wasp per colony/nest for each population (N = 10 nests per population), with the exception of Vero Beach, in which we only found 5 nests. All samples consisted of foundresses (i.e., reproductives) nesting in cabbage palmettos, *Sabal palmetto*. To avoid collecting wasps that may be closely-related, we collected only one nest per palmetto.

### 2.2 Whole Genome Resequencing and Variant Detection

We extracted Genomic DNA from three legs of each wasp using a DNeasy Blood and Tissue kit, following the manufacturer instructions (QIAGEN, Valencia, CA, USA). Paired-end whole genome libraries were prepared by the Cornell University Genomics Facility using the Nextera library kit. Libraries had an average insert size of 550 bp, and were quantified by a bioanalyzer (Agilent Genomics, Santa Clara, CA). We also implemented the recommended Pippin Prep for DNA size selection. Paired-end 150-bp reads were sequenced on 1/6 of a lane of the Illumina NovaSeq 6000. Reads were filtered with Trimmomatic (Bolger et al. 2014) and aligned to our *M. mexicanus* reference genome (Miller et al. 2022) with the Burrows-Wheeler Aligner (v0.7.13) (Li and Durbin 2010). Duplicate sequences were identified with the MarkDuplicates tool within Picard tools (v2.8.2). We called variants with GATK (v3.8) (Van der Auwera et al. 2013), using HaplotypeCaller to first identify variants in each individual, followed by GenotypeGVCF to aggregate variant calls across samples. Further, we applied a hard filter to the combined SNP file to remove poor quality variants using the options Strand bias (FS) > 60, Strand Odds Ratio (SOR) > 3.0), and RMS Mapping Quality (MQ) < 40.0, resulting in a dataset of 2,723,473 SNPs. Given that our system is haplodiploid with high relatedness among siblings, we identified relatives using the *relatedness2* function in VCFtools 276 (v0.1.15) and removed full siblings. Individuals with low average sequencing depth (<1) and high missing data (>35%) were also removed. This resulted in a dataset of 85 individuals with 7-10 samples representing each population, except for the Vero Beach that consisted of 5 samples.

Following our initial filtering, we ran subsequent analyses both on a stringently filtered dataset that did not allow for any missing data per loci, and an imputed dataset that allowed for 20% missingess per loci. Allowing missing data increased the number of SNPs that could be analyzed for adaptive genetic variation from 84,204 to 931,579, and running analyses on both datasets allowed for cross validation of our results. For each dataset, we used VCFtools (v0.1.15) (Danecek et al. 2011) to keep only biallelic SNPs (--max-alleles 2). SNPs with a minor allele frequency less than 0.03 were removed (--maf 0.03), as we would not expect very uncommon alleles to be identified as adaptive loci. To accommodate outlier-detection methods that do not allow for any incomplete data, missing genotypes were imputed using Beagle 5.2 which infers missing SNP data by phasing genotypes (Browning and Browning 2016). The number of iterations was increased from default settings to 30 to improve estimation of genotype phase.

### 2.3 Population structure and effects of geography on genetic variation

We determined broad patterns of genetic variation in our samples by conducting a principal component analysis using Plink’s (v1.9) –pca function (Chang et al. 2015). We then used ADMIXTURE (Alexander et al. 2009) to determine ancestry and the optimal number of genetically distinct population clusters (K) across our latitudinal cline. Each value of K=1-10 was iterated 100 times and the K with the lowest cross-validation error was identified.

To analyze the spatial pattern of genetic variability and population connectivity, we implemented an Estimated Effective Migration Surface (EEMS) approach (Petkova et al. 2016). We modeled estimates of migration rates between locations to identify patterns of gene flow across the landscape. EEMS visualizes deviations from the null expectation of IBD by evaluating the genetic similarity between geographically indexed genetic data. For the 85 re-sequenced genomes, we used common genome-wide SNPs (MAF >5%) and pruned for linkage-disequilibrium (Petkova et al. 2016). We tested a series of deme sizes (100, 200, 400)) with 6 million MCMC iterations and 2 million iterations burnin. Results were checked for MCMC chain convergence and the model fit was evaluated using the linear relation between observed and fitted values for within- and between-deme estimates.

### 2.4 Effects of climate on genetic variation

To identify candidate adaptive loci based on correlations between climatic and genetic data, we used three independent genotype-environment association (GEA) methods that control for population structure and detect signatures of local adaptation (Capblancq et al. 2018, Forester et al. 2018, Duruz et al. 2019, Capblancq and Forester 2021). For all three methods, we used the *raster* R package to implement bioclimatic data for each collecting site from WorldClim at 0.5 (km^2^) resolution. Results from our recent physiological experiments suggest that temperature and precipitation are key variables in thermal tolerance limits. Multicollinearity among climatic variables was assessed by calculating the variance inflation factors to ensure they were less than 5, and that each pair of environmental predictors was correlated less than 0.8 (Forester et al. 2018). Therefore, we focused on four key climatic variables: maximum temperature of the warmest month (BIO5), minimum temperature of the coldest month (BIO6), average precipitation of the wettest month (BIO13) and average precipitation of the driest month (BIO14).

We first used Redundancy Analyses (RDA) to identify covarying allele frequencies associated with the multivariate environmental variables (Capblancq and Forester 2021). We implemented partial RDAs (pRDAs) to decompose the independent effects of climate, neutral genetic variation (e.g., population structure), and geography on genetic variation. This variance partitioning method allowed us to assess the independent contributions of each variable in species that are influenced by IBD and complex demographic histories (Legendre and Legendre 2012, Capblancq and Forester 2021). We then tested and compared the corresponding variance partitioning using either our stringent or imputed genomic datasets. Further, we modeled linear relationships among environmental predictors of interest across our cline, with the *vegan* R package (Forester et al. 2018, Capblancq and Forester 2021). Significance for each RDA model was tested using 999 permutations of an ANOVA-like test with the built-in function “anova.cca()”. All axes were significant at 0.01. Identified outlier SNPs had a ± 3 *SD* from the mean score of each constrained axis and a locus score of ±2.5 (Forester et al. 2018).

Second, we performed outlier SNP detection using latent factor mixed models (LFMM) with the *LEA* R package (Caye et al. 2019). The number of latent factors was identified as K= 2 based on the ADMIXTURE results, as well as *LEA*’s built-in “snmf()” function, which employs sparse Non-Negative Matrix Factorization algorithms. LFMM analyses require no missing genomic data, requiring us to impute any missing genotypes. We performed a Bonferroni correction on the p-values (<0.05), which resulted in a conservative set of outliers.

Finally, we identified outlier SNPs using the *R.SamBada* R package, designed to search for signatures of local adaptation (Duruz et al. 2019). This method computes logistic regression models to test the relationship between a binary genetic variable (i.e., alternative alleles) and environmental predictors. We first tested for the same correlation matrix for the four bioclimatic predictors with r^2^ < 80% by using the *prepareEnv* function, as recommended to validate their independence (Duruz et al. 2019). We then implemented *SNPRelate* to run a principal component analysis (PCA) to assess the genetic structure of each population. To be consistent with our other GEA methods, we also used K=2 to inform the model and a Bonferroni correction of p-values < 0.05 as a cutoff for significance. Given the two genetic clusters shown by the ADMIXTURE analysis, we ran R.SamBada in bivariate mode by adding the score of the first PCA to account for population structure (as in Duruz et al. 2019).

To identify candidate SNPs associated with our four bioclimatic predictors (BIO5, BIO6, BIO13 and BIO14), we only included SNPs that were consistently significant for all three GEA methods, using the conservative cutoff of Bonferroni-corrected p-values. These outlier SNPs were matched to their associated genes using the annotated genome for *Mischocyttarus mexicanus* (JACGTP000000000). SNPs that occurred in regions that were not annotated were further investigated using NCBI BLAST. Gene regions were viewed using the Integrative Genomics Viewer (IGV) and the surrounding sequences were extracted. BLASTn searches were conducted systematically against the arthropod and hymenopteran databases, in addition to available *Polistes, Apis,* and *Drosophila* genomes. Annotated genes with E-values less than 1e-50 were then BLASTed back to the *Mischocyttarus mexicanus* genome for validation.

Synteny of gene regions in related *Polistes* genomes were checked to increase confidence. The location of outlier SNPs within the genes was determined using transcriptomic data included in the *Mischocyttarus mexicanus* annotation when available, or through the alignment of the gene region to closely related *Polistes* transcript annotations. Given that the SamBada method can reconstruct the spatial distribution of specific genotypes, we also explored changes in allele frequency across our transect for three consistent and robust candidate SNPs linked to BIO6 (minimum temperature during coldest month) and BIO13 (average precipitation during the wettest month), the two strongest bioclimatic predictors.

## 3 RESULTS

### 3.1 Population structure and effects of geography on genetic variation

We used three approaches to estimate population structure and patterns of genetic differentiation associated with geography. First, ADMIXTURE analysis found that populations of *M. mexicanus* across the transect grouped into two genetic clusters (K =2), which represented the lowest cross-validation error compared to clusters of 3 or 4) (Fig. 1B, Supplementary Figure 1). This pattern shows clear separation of the southern (subtropical) and northern (temperate) populations (Fig 1). In addition, we identified three populations with mixed ancestries (Fig. 1B). Two of these populations, Vero Beach and Orlando, coincide with the transition zone from subtropical to temperate populations in Central Florida. Surprisingly, we also discovered that two individuals from the northernmost population in Charleston grouped with the southern genetic cluster. Second, PC1 for genetic variation (14%) showed a strong correlation with latitude based on the imputed dataset (Spearman rank correlation, *r* = −0.792, *p* < 0.001, N = 85, Fig 1C). Similarly, PC1 based on the stringent dataset explained 16% of the genetic variation and was correlated with latitude (Spearman rank correlation, *r* = −0.786, *p* < 0.001, N = 85). These results match both the gradient in minimum temperature when plotting BIO6 (minimum temperature of the coldest month) across our transect and the Köppen-Geiger climate classification. We obtained the same result for the stringent and imputed datasets, which provides confidence in our imputed genotypes. Third, the IBD analysis showed a significant correlation between linearized genetic distance and geographic distance (y=0.00017x-0.47, R^2^=0.78, P< 0.001, Supplementary Figure 2) for both the stringent and imputed datasets.

Levels of gene flow can be inferred from extent of population structure between adjacent populations. However, we can also more explicitly consider gene flow while controlling for the effects of isolation by distance. The EEMS analysis showed that gene flow between some adjacent populations is reduced, failing to match those expected by a pure IBD model (Supplementary Figure 3).

### 3.2 Effect of climate on genetic variation

We used variance partitioning to disentangle the contributions of population structure, geography and climate on genetic variation across the transect, by implementing pRDA models. Combined, all three factors in our full model accounted for 17.4% of the total genetic variation across our 10 wasp populations (P < 0.001, for statistics of the full model see Table 1). The sole effects of geography and genetic structure on genetic variation were 3.6% (21% of the explained variation) and 2.2% (12.6% of the explained variation), respectively. After controlling for population structure and geography, the effect of climate was significantly higher, accounting for 7.2% of the total genetic variance (41.5% of the explained variation). The pRDA models based on the imputed dataset (Table 1) and the stringent dataset with no missing data provide similar results (Supplementary Table 1).

**Table 1.** Contributions of local climate and geography to genomic variation decomposed by a partial redundancy analysis (pRDA) . The influence of climate is larger than both the influence of population structure and geography on genetic variation for both the imputed and stringent datasets. The proportion of total variance, proportion of explained variance, R^2^, and its significance level are shown. The shown results correspond to the imputed dataset. For pRDA models from the stringent dataset that show the same patterns, see Supplementary Table 1.

| Partial RDA models | Inertia | $R^2$ | p (>F) | Proportion of explainable variance | Proportion of total Variance |
| --- | --- | --- | --- | --- | --- |
| Full model: $F \sim \text{clim.} + \text{geog.} + \text{struct.}$ | 161464 | 0.174 | 0.001 | 1 | 0.17 |
| Pure climate: $F \sim \text{clim.} \mid (\text{geog.} + \text{struct.})$ | 67022 | 0.072 | 0.001 | 0.415 | 0.07 |
| Pure structure: $F \sim \text{struct.} \mid (\text{clim.} + \text{geog.})$ | 20397 | 0.022 | 0.001 | 0.126 | 0.02 |
| Pure geography: $F \sim \text{geog.} \mid (\text{clim.} + \text{struct.})$ | 33339 | 0.036 | 0.001 | 0.207 | 0.04 |
| Confounded climate/structure/geography | 40706 |  |  |  | 0.04 |
| Total unexplained | 768953 |  |  |  | 0.83 |
| Total inertia | 930417 |  |  |  | 1.00 |

The RDA analysis detected 6,535 outlier SNPs associated with maximum temperature (BIO5), minimum temperature (BIO6), and maximum precipitation (BIO14). There were no outlier SNPs associated with minimum precipitation (BIO 13). RDA1 accounted for 49.7% of the variation and separated northern and southern populations, with Vero Beach as the intermediate population (Fig 2A). RDA2 accounted for 19.1% of the variation and further separated the northern and southern individuals into their respective populations. The only two samples that did not group geographically were the two Charleston outliers, which was expected due to their shared genomic background with southern populations. The inset map reflects the total SNPs as a gray block plotted along the same RDA axes, visualizing outlier SNPs as the points farthest from the center (Fig 2b). However, because most of 6,535 outlier SNPs showed a weak association with the four bioclimatic predictors, we only classified candidate SNPs as those that were also detected by the other two GEA methods for subsequent analyses.

**Fig 2.**
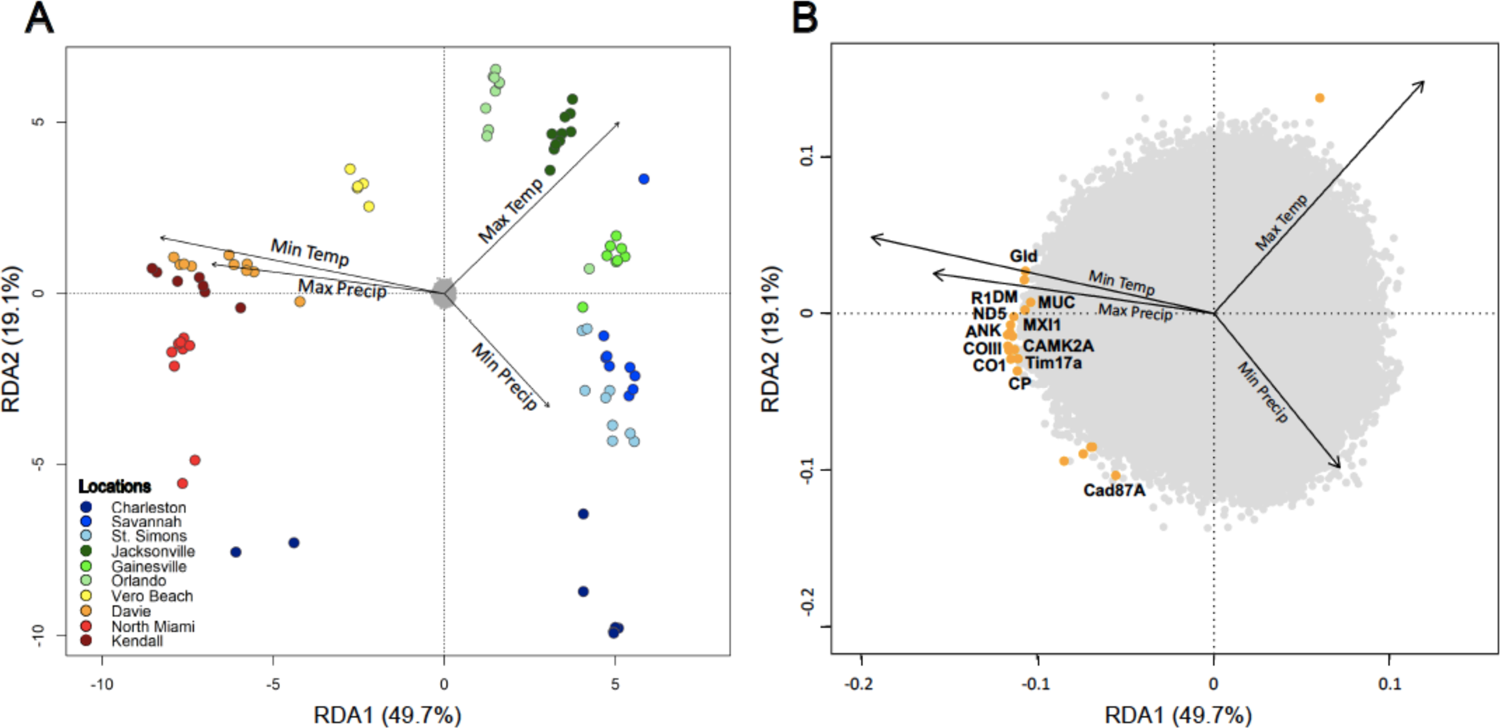
Redundancy (RDA) results of the genotype-environment association. **A.** Orthogonal projections of the first two RDA axes represent 69% of the explained variance. Arrows correspond to the four bioclimatic predictors that were significantly associated to specific populations across the climatic transect (P <0.05). Arrow length represents the strength of the relationship with individual axes, while their directions indicate positively and negatively correlated variables in relations to distinct populations. Wasp samples are color coded by their population of origin and oriented with the positioning of single nucleotide polymorphisms (SNPs) as a gray block in the center of the plot. **B.** Plot shows a zoom-in of the SNPs block, highlighting SNPs with a significant association (orange dots) to the bioclimatic predictors. Outlier SNPs are positioned on the outer edge of the block (± 3 SD) while SNPs that show weaker or no association with the environmental variables are inside the block (gray dots).

The more conservative LFMM analysis found a total of 36 outlier SNPs, but the majority of these highly significant outliers are the same as those detected by the RDA analyses. For instance, the specific variants with *q*-values < 0.05 showed the same association patterns with bioclimatic variables, cross-validating the two GEA methods (Fig 2B, Fig 3). Finally, the R.SamBada identified 1899 SNPs, after Bonferroni-correction(P <0.05). In sum, 24 outlier SNPs were consistently detected by the three GEA analyses (Fig 4, Table 1). The majority of these 24 shared outliers were associated with minimum temperature, followed by maximum precipitation. Two of these 24 shared outlier SNPs were associated with maximum temperature. (Fig 2B and Fig 3).

**Fig 3.**
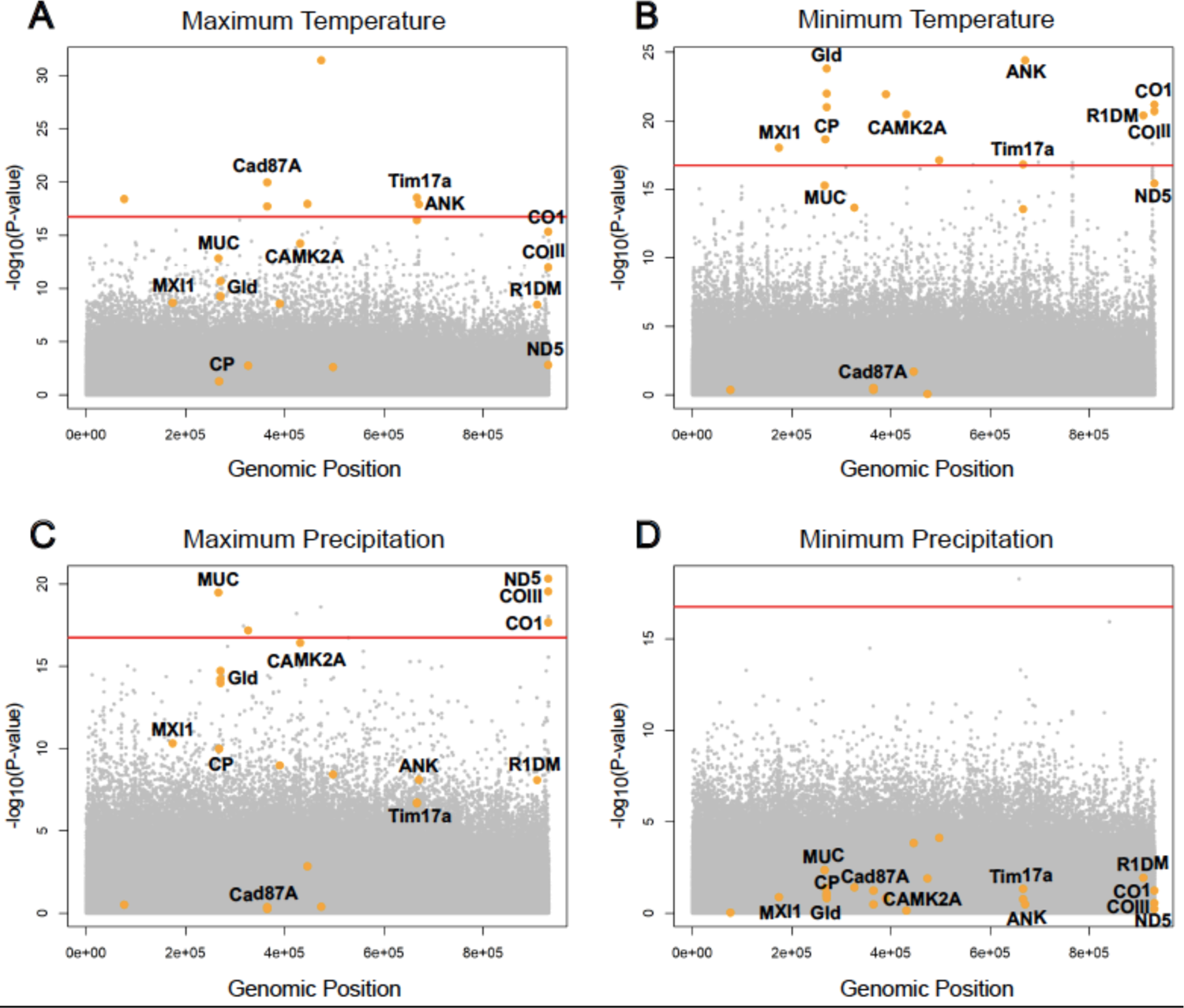
Manhattan plots of single nucleotide polymorphisms (SNPs) that were consistently associated with each four bioclimatic predictors using the RDA, LMFF and SamBada genotype-association approaches (GEA). SNPs of interest, given their significant associations to one or more bioclimatic variables, are represented by orange circles for A) maximum temperature for the warmest month (BIO5), B) minimum temperature of the coldest month (BIO6), C) precipitation of wettest month (BIO13) and D) precipitation of driest month (BIO14). The y axis represents the -log_10_(P) values, with P being the probability of association between a bioclimatic variable and a SNP. The red line depicts the 10^−5^ *p*-value threshold. The outlier SNPs above the significance threshold are consistent with the majority of the outlier SNPs found in the RDA analyses.

**Fig 4.**
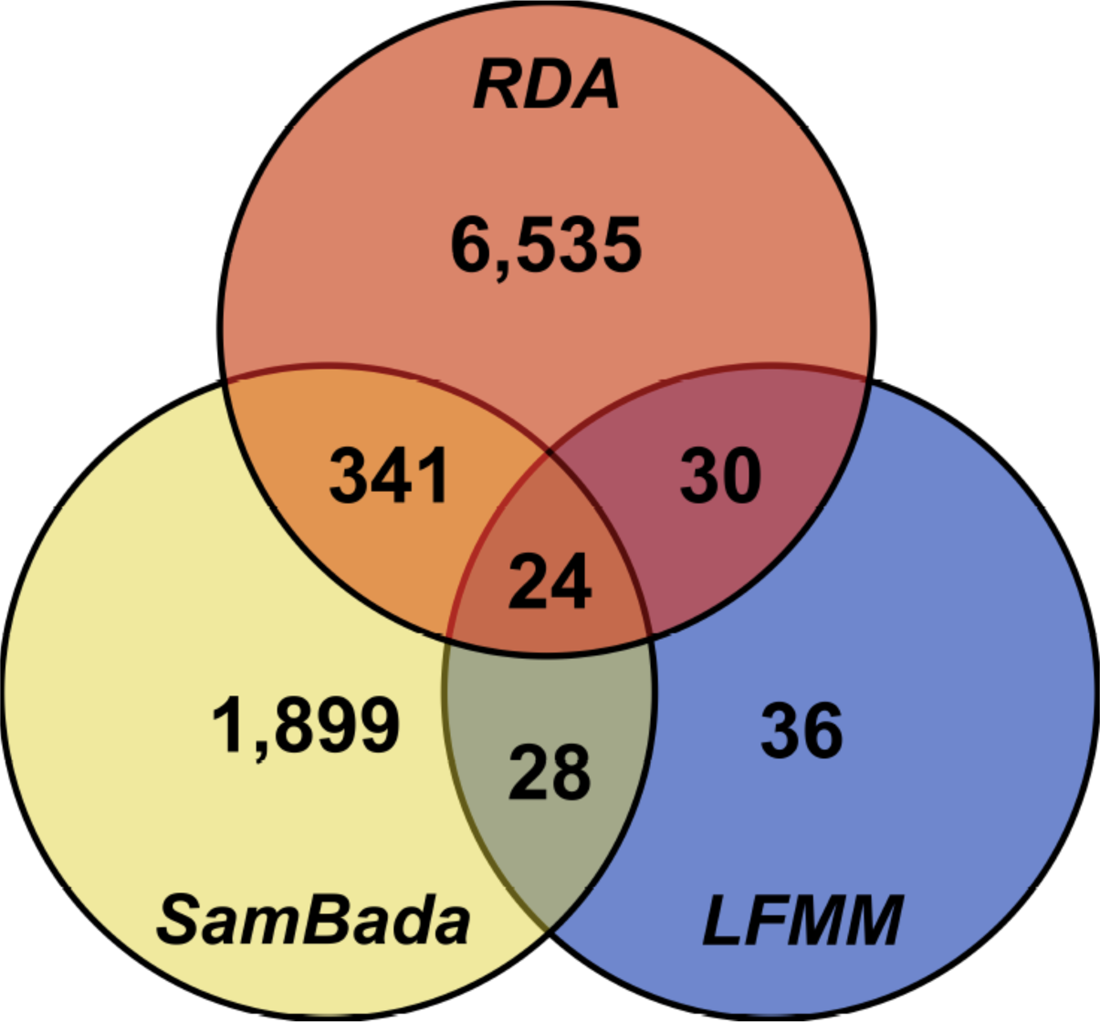
Venn diagram depicting the number of significant SNPs consistently detected by the three GEA methods. The redundancy analysis (RDA) is the most liberal method, detecting many SNPs with weak associations, while the latent factor mixed model (LFMM) is the most conservative method, implementing a stringent Bonferroni correction. We identified a total of 24 candidate outlier SNPs that consistently showed to be under selection for all three GEA approaches.

### 3.3 Candidate SNPs suggest local adaptation

The signatures of local adaptation detected by all three GEA approaches revealed 24 SNPs that are associated with changes in climatic variables across the transect. Annotation against the *M. mexicanus* genome, along with confirmation against three available *Polistes* genomes, revealed known genes and their functions for 19 of the 24 candidate SNPs (Table 2). A total of 16 SNPs were associated with BIO6 near or withing genes known to affect thermal tolerance and/or mitochondrial metabolism (Table 1). Three SNPs from the glucose dehydrogenase gene (*Gld*) were consistently linked with minimum temperature. Two of these *Gld* SNPs showed non-synonymous mutations. Single SNPs in the *CO1* and *COIII* mitochondrial genes were also associated with minimum temperature. A SNP in another mitochondrial gene, NADH dehydrogenase subunit 5 (*ND5)*, was linked with both minimum temperature and maximum precipitation. A SNP in the max-interacting protein 1-like(MXI1) gene, implicated in bioenergetics, was also associated with minimum temperature. The allelic variants of the *Gld*, *ND5* and *MXI1* SNPs showed a spatial distribution that matched the climatic gradient (Fig. 5).

**Fig 5.**
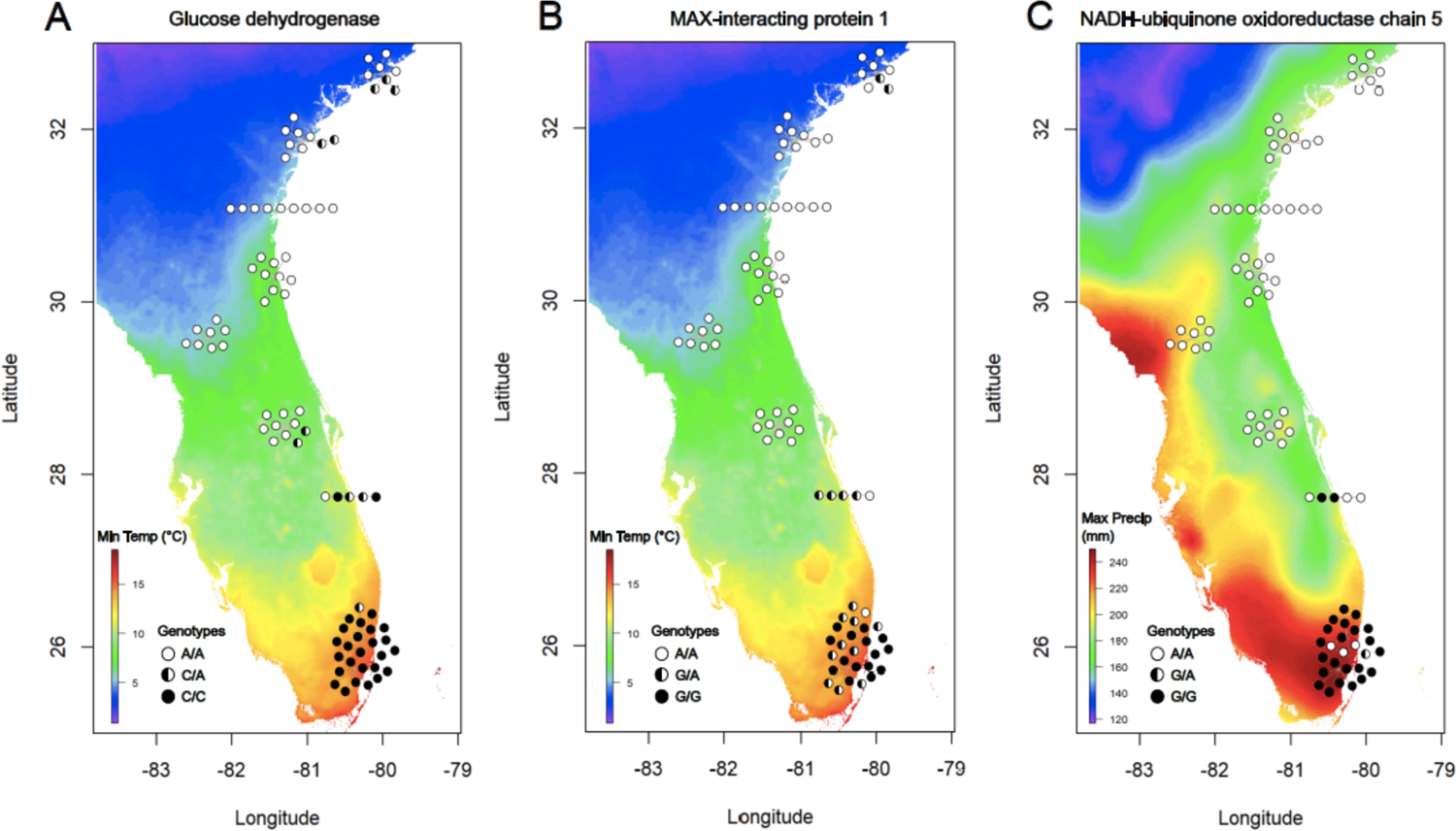
Spatial distribution of genotypes for SNPs associated with bioclimatic predictors. For each map, the color of the background shows changes in the bioclimatic predictor across the transect. Color-coded circles indicate the distribution of individuals of a specific genotype under climatic selection. **A.** Glucose dehydrogenase SNP (*GLD.1605462*) and **B.** Max-interacting protein 1-like SNP (*MXI-1*) were both associated with the average minimum temperature of the coldest month (BIO6). **C.** NADH dehydrogenage subunit 5 SNP (*ND5*) was associated with maximum average precipitation (BIO5). All three SNPs showed the same pattern, indicating a prevalent genotype in the subtropical populations and a different genotype prevalent in temperate populations. Heterozygotes and/or a combination of genotypes are most common in the transition zone.

**Table 2.**
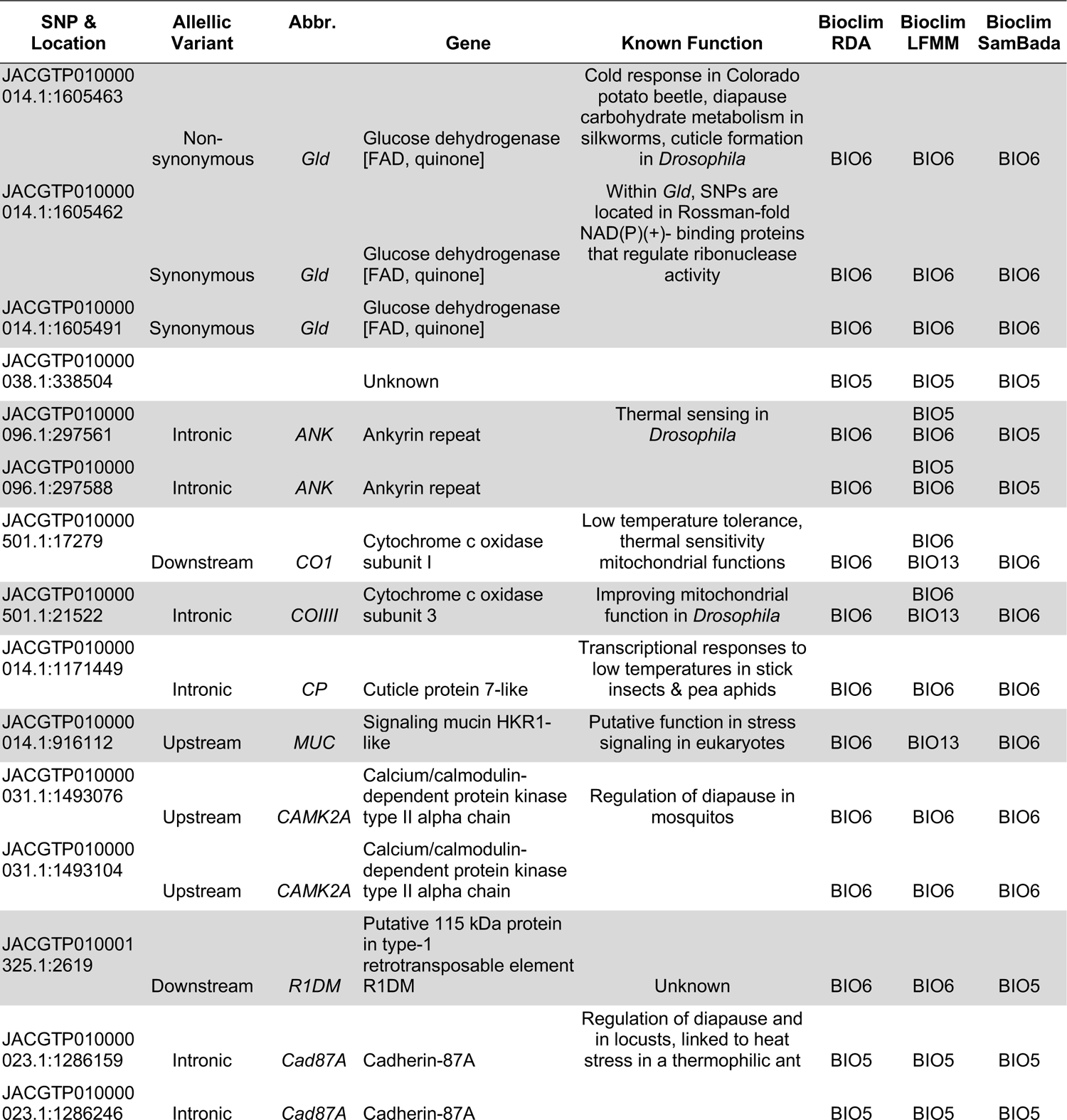
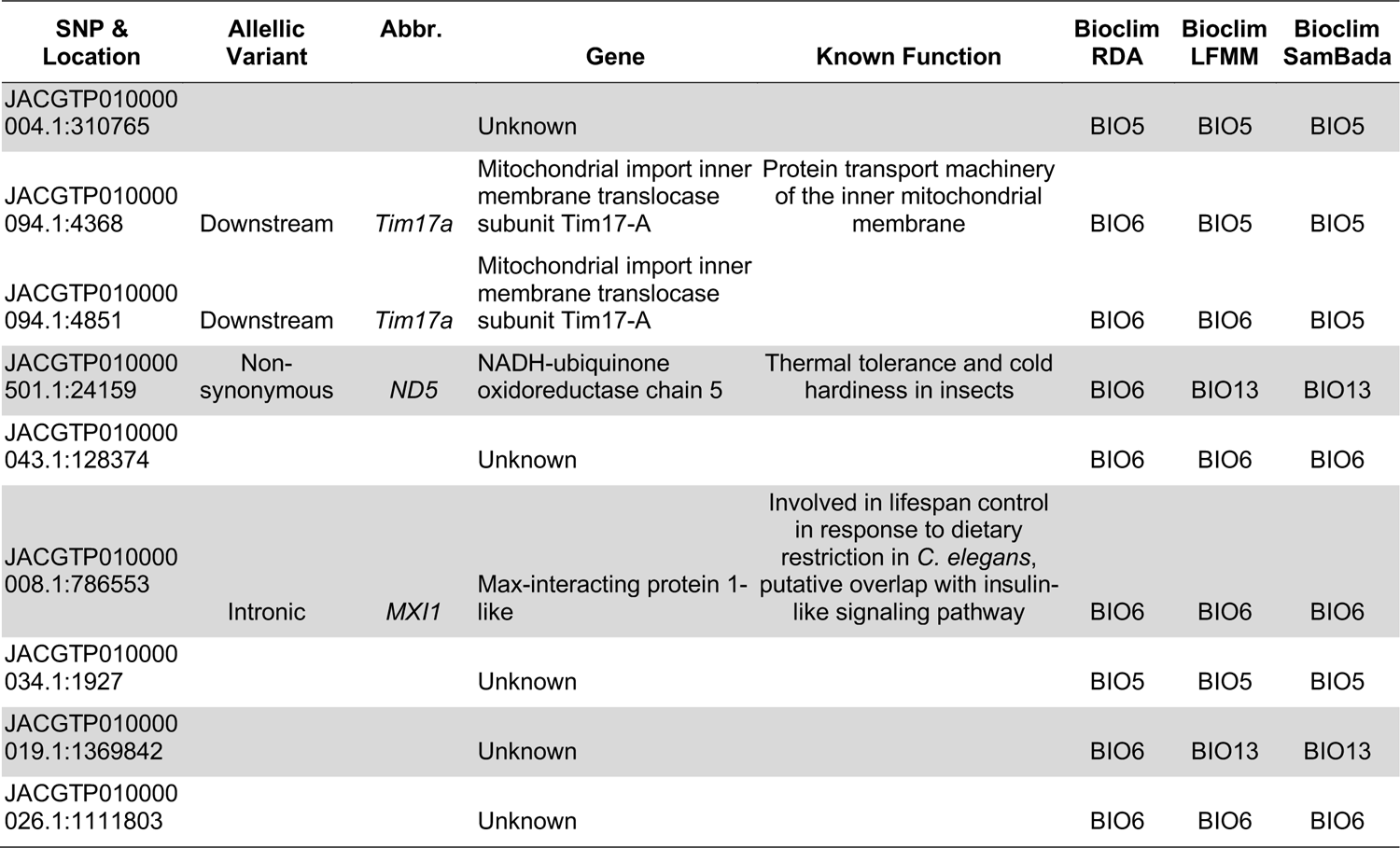
Outlier SNPs identified with annotations and putative functions, validated by all three GEA approaches independently.

The two outlier wasps from the northernmost population that grouped with the southern genetic cluster also showed temperate genotypes for these same SNPs (Fig. 5).

## DISCUSSION

In this study, we aimed to identify the main drivers of local adaptation in natural populations of the wasp *Mischocyttarus mexicanus*, a species with a wide distribution across a latitudinal and climatic gradient. We found strong population structure along with an effect of isolation by distance across the range similar to a previous study in another paper wasp, *Polistes fuscatus* (Bluher et al. 2020). However, patterns of gene flow along the range were variable, with some adjacent regions showing reduced gene flow compared to that expected by IBD alone. In these cases, gene flow may be reduced due to transition in climate (i.e., subtropical to temperate) or fragmentation caused by differences in habitat quality and land use. Importantly, the three independent GEA analyses revealed a stronger effect of climate than population structure and geography in explaining genetic variation across the transect. We found evidence that outlier SNPs may be under selection and facilitated local adaptation across this climatic transect. For instance, several SNPs are strongly associated with the transition from subtropical populations that have year-round active nests to temperate zone populations that experience seasonality and have adapted to overwinter. In particular, the majority of the outlier SNPs were near or at genes that are associated with thermal tolerance, and a few were linked to maximum precipitation across the gradient. These results are consistent with previous studies in wild populations of *Drosophila melanogaster* that show climatic and seasonal adaptations across Europe and North America (Telonis-Scott et al. 2011, Mateo et al. 2018, Bogaerts-Márquez et al. 2021, Machado et al. 2021). Our findings are also consistent with population-specific signatures of local adaptation to thermal tolerance discovered in bumblebees (Jackson et al. 2020, Pimsler et al. 2020, Heraghty et al. 2022) and solitary bees (Samad-zada et al. 2023) with similarly broad distributions.

Compellingly, the detected links between climate and genetic variation mirror our recent thermal tolerance experiments in wasps collected across the same climatic transect. In northern populations, reproductive wasps that overwinter and initiate nests the following spring show a higher tolerance to extended cold temperatures compared to workers that only live during the late spring and summer (Baudier, Robinson, Uy, *unpublished data)*. Further, subtropical populations that experience high temperatures most of the year show a higher critical thermal maximum and survival to extended high temperatures compared to seasonal northern populations (Robinson, Uy, Baudier *unpublished data*). Together, the genetic variation patterns, thermal tolerance and nest initiation strategies suggest that specific genes facilitate local adaptation for *M. mexicanus,* allowing it to expand its range and adapt to the temperate zone’s seasonality constraints.

Overall, outlier SNPs showed a strong association with the minimum temperature of the coldest month, followed by maximum precipitation and maximum temperature. Several outlier SNPs were, in turn, at or near genes known to mediate thermal tolerance. One particular candidate gene of interest is glucose dehydrogenase (*Gld)*, which catalyzes the oxidation of glucose to gluconolactone and is involved in cuticle formation in *Drosophila* (Cavener et al. 1986), as well as cold tolerance responses in beetles (Govaere et al. 2019) and diapause in *Bombyx* silk worms (Jiang et al. 2019). Additionally, three different SNPs in *Gld* are associated with minimum winter temperature. We hypothesize that *Gld* is critical for temperate populations that experience colder winters and relatively less rainfall than the subtropical populations, as desiccation stress may be more prevalent and cuticle formation may be particularly relevant. Two of the *Gld* allelic variants result in non-synonymous mutations unique to the expansion into temperate zones, suggesting selection on these variants for enhanced survival in cold temperatures.

Another candidate gene we detected mediates the production of NADH dehydrogenase subunit 5 (ND5), a mitochondrial enzyme that is a core subunit of Complex I in the electron transport chain. The mitochondrion is a hub of energy division for insects experiencing cold stress (Lubawy et al. 2022). Notably, the frequencies of allelic variants for *ND5* also show a distinct pattern of spatial distribution following minimum temperature transitions across the climatic cline (see Fig. 5). Intriguingly, two other outlier SNPs occur in the *CO1* and *COIII* genes, respectively. Both genes are involved in the mitochondrial electron transport chain, showing transcriptional responses to low temperature tolerance in insects (Huang et al. 2017, Noer et al. 2023). Overall, our results support the important role of the mitochondria as a center for genes that may be under strong climate-driven selection, resulting in locally adapted populations.

Another outlier SNP is found in an intronic region of *MXI-1, a gene* associated with lifespan and energetic needs (Johnson et al. 2014), which suggests a potential regulatory function in metabolism. The *MXI-1* SNP shows a similar pattern of climatic transition in the spatial distribution of genotypes as the other candidate SNPs. Remarkably, although the two wasps from the northernmost population grouped with the subtropical genomic cluster, they were either heterozygous or homozygous for the temperate allelic variants for *Gld, ND5* and *MXI-1*. These results suggests that these individuals have the right allelic variants to adapt to temperate zones, likely due to offspring of long-distance dispersers mating with a northern population. Finally, several of the other outlier SNPs fall within intronic regions and one SNP with unknown function is located in a transposable element. The effects of these SNPs require further exploration, as other studies implicate regulatory regions and transposable elements underlying stress responses elicited by harsh environmental conditions in natural populations of *Drosophila* (Casacuberta and González 2013, Lidia et al. 2013, Lerat et al. 2019, Green et al. 2022). Future gene expression and functional testing experiments will confirm the role of these candidate SNPs in facilitating local adaptation.

## Conclusions

To predict the response of species to impending anthropogenic climate change, it is necessary to infer if they can adapt to rapid changes (Aitken and Whitlock 2013). The ability of *M. mexicanus,* and other species with a broad distribution, to adapt to climate change depends on both phenotypic plasticity and the genetic diversity available to selection (Menzel and Feldmeyer 2021, Vranken et al. 2021, Martin et al. 2023). *Mischocyttarus* is a neotropical genus, with only a few species that expanded to temperate zones (Silveira 2008, Somavilla et al. 2021). Our work suggests a northward expansion that likely depended on genomic evolution that facilitated adaptive responses to seasonality, cold tolerance, and desiccation. However, with unprecedented rising temperatures, the ability of *M. mexicanus* to persist in its current range or expand further north remains unclear (Menzel and Feldmeyer 2021). As global climate change rapidly advances, social insects that provide ecosystem services such as predation and pollinations may be highly affected. The integration of genomic and environmental information can be a powerful approach to predict the vulnerability and persistence of species under climate change.

## AUTHOR CONTRIBUTIONS

This study was conceived by FMKU with input from SEM and MJS. FMKU collected field specimens along the transect and extracted DNA for sequencing. HLC, GG, SEM, KL and FMKU performed analyses. HLC and FMKU wrote the manuscript with input from SEM. All authors reviewed, edited and approved the final manuscript.

## ACKNOWLEDGEMENTS

We thank members of the community for letting us collect wasps in their properties across the South East USA. Clark Alexander and the Skidaway Institute of Oceanography, University of Georgia, provided collecting and logistical support. HLC, GG and FMKU were supported by funding provided by the University of Rochester. MJS and SEM were supported by NSF DEB CAREER 1750394 awarded to MJS. Al Uy, members of the Uy lab and Christopher Jernigan provided thoughtful comments in previous versions of the manuscript.

