## Supplementary Material for "Adaptive genomic variation is linked to a climatic gradient in a social wasp"

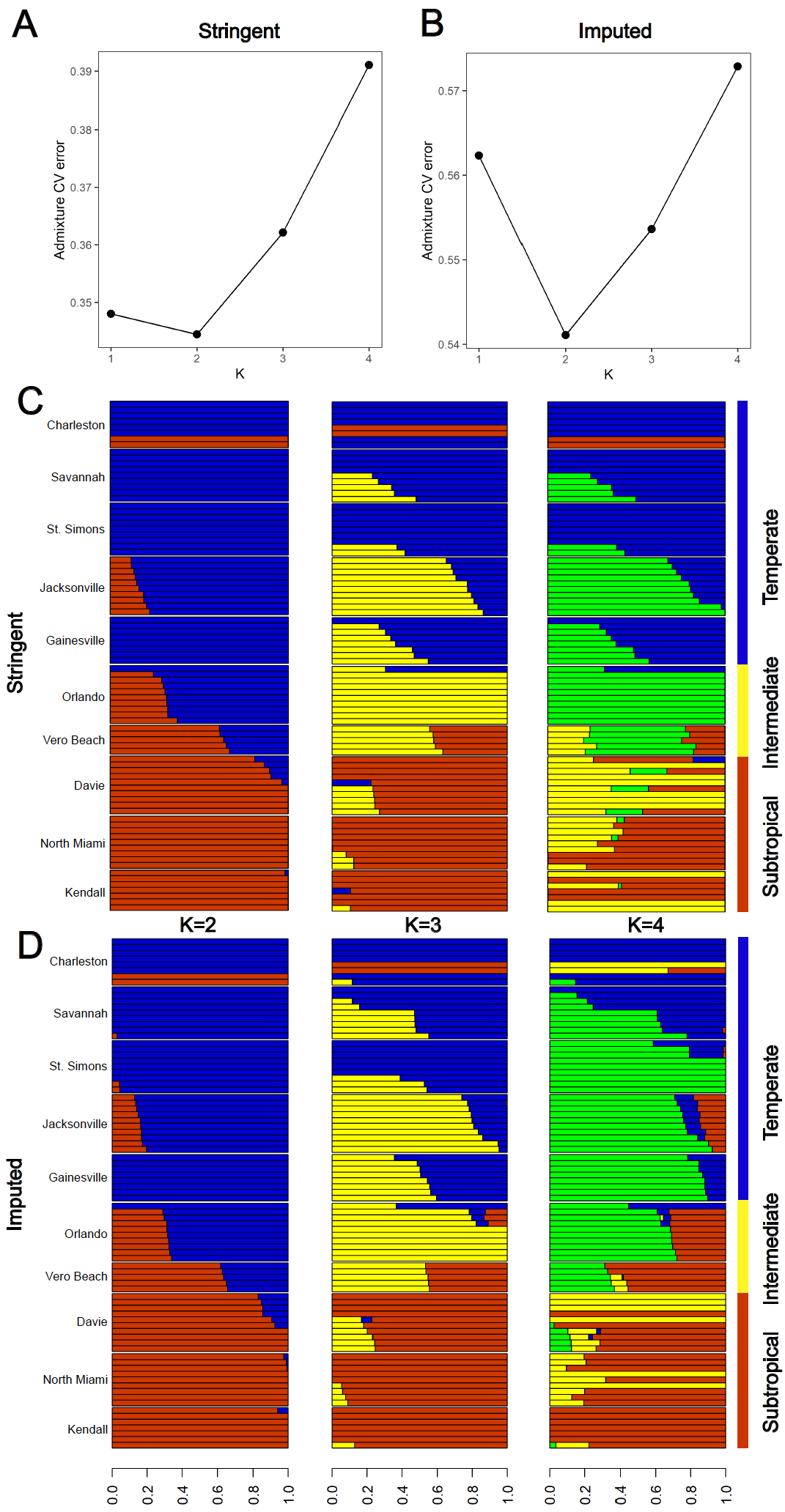

**Supplementary Figure 1**. **ADMIXTURE analysis of *Mischocyttarus mexicanus* populations.** Mean likelihood values for delta K, with K=2 showing the best fit likelihood value based on the **A**) imputed and **B)** stringent datasets. **C.** Structure plots from K=2 to K=5 for the ten populations across the latitudinal and climatic gradient .

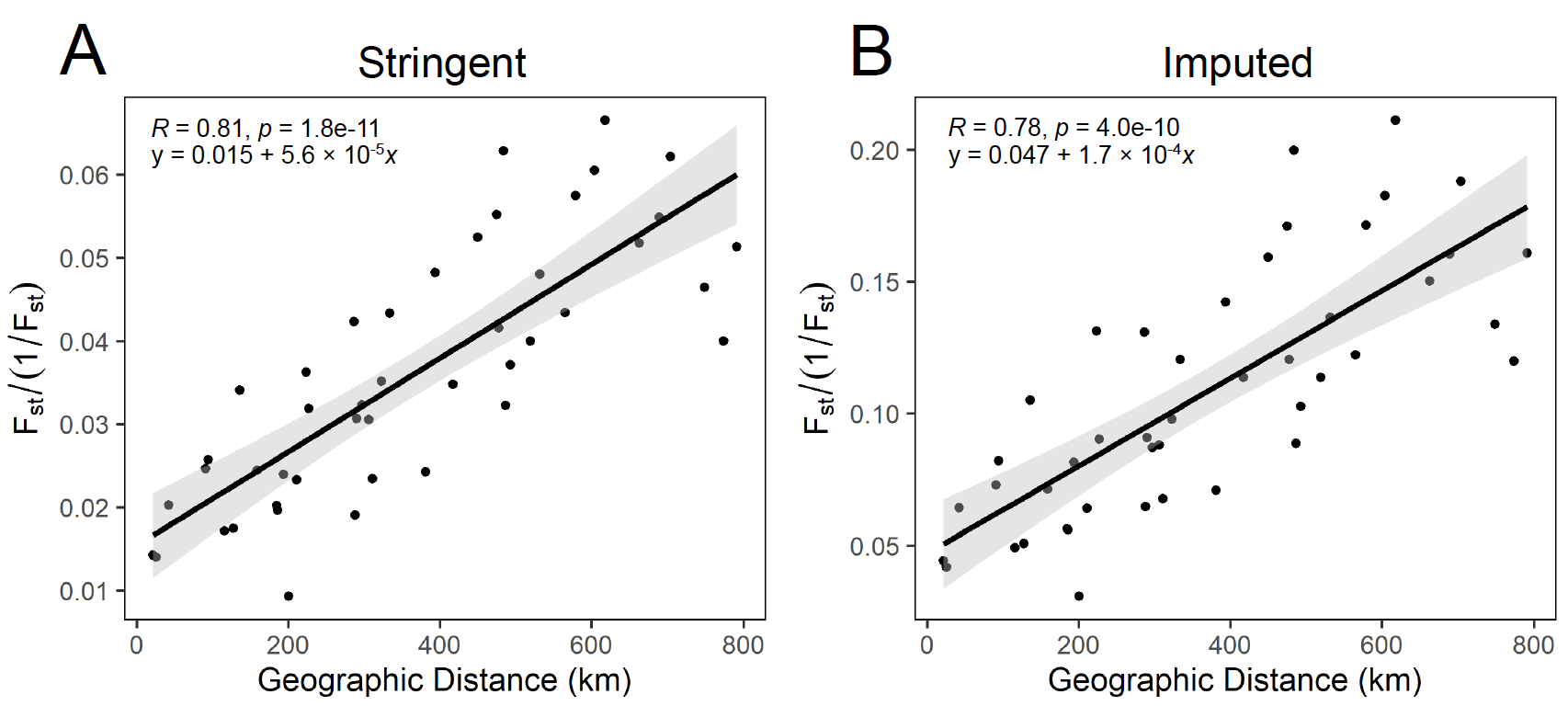

**Supplementary Figure 2**. **Patterns of isolation by distance for M. mexicanus across the Southeastern transect.** Strong positive correlations between geographic distance and genetic distance for both the A) original dataset and B) the imputed dataset.

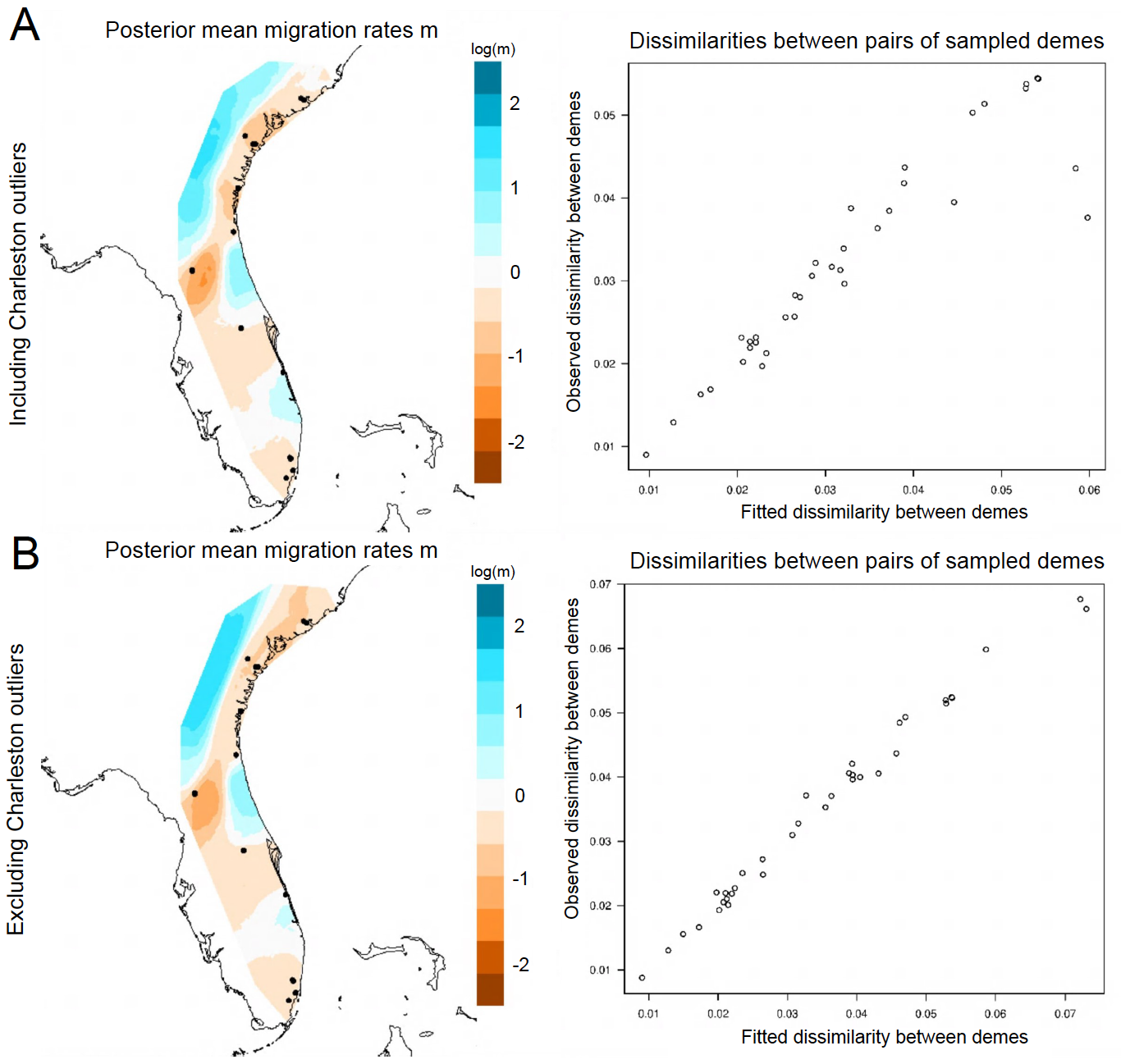

**Supplementary Figure 3**. Visualizing spatial population structure of *M. mexicanus* using estimated effective migration surfaces (EEMS). Samples were collected across the latitudinal transect that shows a climatic gradient. The size of the dots indicated the number of individuals sequenced per population and the scale indicates patterns of gene flow, with orange representing reduced gene flow. Gene flow patterns do not change when A) including and B) excluding the two samples from Charleston that group with the southern genetic cluster.

**Supplementary Table 1.** **The contribution of local climate towards genetic variation is larger compared to the contributions of geography and population structure.** Results from a partial redundancy analysis (pRDA) depict the proportion of total variance, proportion of explained variance, R^2^, and its significance levels. The analysis is based on the stringent dataset with no missing values. For consistent results from the imputed dataset, see Table 1.

| **Partial RDA models** | **Inertia** | **R^2^** | **p (>F)** | **Proportion of explainable variance** | **Proportion of total Variance** |
| --- | --- | --- | --- | --- | --- |
| Full model: F ~ clim. + geog. + struct. | 11891 | 0.143 | 0.001 | 1 | 0.14 |
| Pure climate: F ~ clim. \| (geog. + struct.) | 5265 | 0.063 | 0.001 | 0.443 | 0.06 |
| Pure structure: F ~ struct. \| (clim. + geog.) | 1527 | 0.009 | 0.001 | 0.128 | 0.02 |
| Pure geography: F ~ geog. \| (clim. + struct.) | 2552 | 0.031 | 0.001 | 0.215 | 0.03 |
| Confounded climate/structure/geography | 2547 |  |  |  | 0.03 |
| Total unexplained | 71293 |  |  |  | 0.86 |
| Total inertia | 83184 |  |  |  | 1.00 |
